# Automated cell annotation in scRNA-seq data using unique marker gene sets

**DOI:** 10.1101/2024.05.24.595477

**Authors:** Linh Truong, Thao Truong, Huy Nguyen

## Abstract

Single-cell RNA sequencing has revolutionized the study of cellular heterogeneity, yet accurate cell type annotation remains a significant challenge. Inconsistent labels, technological variability, and limitations in transferring annotations from reference datasets hinder precise annotation. This study presents a novel approach for accurate cell type annotation in scRNA-seq data using unique marker gene sets. By manually curating cell type names and markers from 280 publications, we verified marker expression profiles across these datasets and unified the nomenclature to consistently identify 166 cell types and subtypes. Our customized algorithm, which builds on the AUCell method, achieves accurate cell labeling at single-cell resolution and surpasses the performance of reference-based tools like Azimuth, especially in distinguishing closely related subtypes. To enhance accessibility and practical utility for researchers, we have also developed a user-friendly application that automates the cell typing process, enabling efficient verification and supporting comprehensive downstream analyses.

## Introduction

The advent of single-cell RNA sequencing (scRNA-seq) technology has revolutionized the field of biology by enabling the transcriptome-wide analysis of individual cells. This powerful tool has provided unprecedented insights into the complexity and heterogeneity of biological systems, allowing researchers to characterize cellular diversity, identify rare cell populations, and elucidate cellular differentiation trajectories. However, the analysis of scRNA-seq data requires accurate cell type annotation, which remains a significant challenge. This process is typically manual, requiring researchers to painstakingly verify marker genes across all known cell types and subtypes. This is highly time-consuming and error-prone due to several factors. Primary among these is the inconsistency in cell type definitions across different studies. Researchers often use varied nomenclature and classification criteria, which can lead to discrepancies in cell type identification and complicate comparative analyses. Moreover, the continuous discovery of new cell types and subtypes, along with the lack of standardized nomenclature, further complicates the accurate annotation of cell types. Technological variability further complicates cell type annotation. Differences in scRNA-seq platforms and protocols can affect gene expression profiling, leading to variability in data that can misguide annotation efforts if not properly adjusted. While labeling well-separated clusters of large cell populations is manageable, many subtypes are closely related and reside within the same cluster (Figure 1A). This makes separating them difficult, especially when batch effects mix them further together (Figure 1B) or when the clustering does not align with the marker expression pattern (Figure 1C).

**Figure 1.**
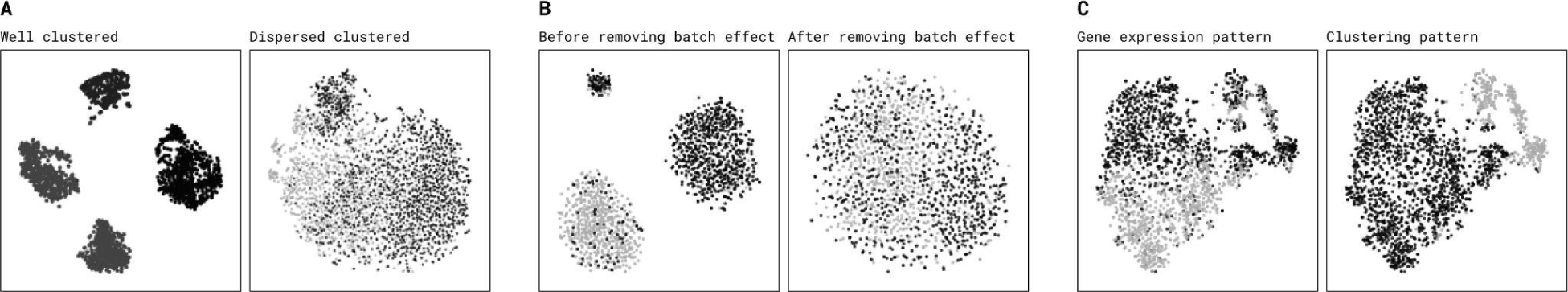
Challenges in cell type annotation using clustering methods. **A.** Broad cell types with distinct expression profiles are usually well-separated on t-SNE or UMAP visualizations, allowing for easy identification either through clustering or manual selection. In contrast, cell subtypes often distribute closely and overlap, making manual separation impractical. **B.** The batch correction process tends to homogenize the differences between cell subtypes, resulting in a mixed population that cannot be distinguished by clustering or manual methods. **C.** While closely related cell subtypes can be differentiated using canonical markers, they are often indistinguishable when relying solely on clustering patterns.

To address these challenges, several computational methods have been developed, including reference-based mapping tools such as Azimuth^3^. These tools aim to expedite the annotation process by transferring labels from annotated reference datasets to new ones based on cell similarity. However, such methods still have limitations that can’t fully replace manual curation. Differences in technological platforms between query and reference datasets often prevent accurate mapping, resulting in poor agreement with original annotations even when querying a reference against itself. Additionally, differences in cell type compositions between the query and reference datasets can lead to misaligned cell type labels, forcing researchers to revert to manual curation. Lastly, reference-based mapping relies heavily on clustering methods, which often fail to distinguish closely related subtypes like immune cells. Therefore, there is an urgent need for a more robust and accurate approach to cell type annotation in scRNA-seq data.

In this study, we present a novel method that utilizes unique marker gene sets, derived from a rigorous curation of 280 publications, to consistently identify cell types and subtypes. Our approach addresses the aforementioned challenges and provides a reliable solution for accurate cell type annotation in scRNA-seq data. The customized algorithm based on the AUCell method accurately labels cells at single-cell resolution and surpasses the performance of the Azimuth tool, particularly in identifying closely related subtypes. Furthermore, we have developed a user-friendly application that automates the cell typing process, enabling efficient verification and supporting comprehensive downstream analyses. The application is accessible at https://omnibusx.com. Our method simplifies the cell type annotation process, eliminates the need for extensive reference selection, and is capable of identifying both broad and narrowly defined cell populations that might be overlooked by other methods.

## Results

### Curate comprehensive marker gene sets

To address the challenges of cell type annotation in scRNA-seq data, we focused on building a comprehensive and accurate map of cell type and subtype-specific marker genes. Over the past decade, extensive research has been conducted to study and verify these markers. However, inconsistencies in naming conventions across the scientific community and a lack of cross-study validation have led to ambiguous and non-reproducible marker gene sets. To overcome these challenges, we undertook a detailed and rigorous curation process, involving multiple steps to refine and validate marker gene sets from 280 publications (Supplementary Data 1), ensuring their specificity and utility in accurately annotating cell types across various studies and tissues.

Our first step was to conduct a comprehensive literature review to collect existing knowledge on marker genes associated with different cell types and subtypes. This extensive collection phase aimed to compile a preliminary list of markers that have been validated and used in previous studies. We then enhanced these marker lists by performing t-tests to extract lists of differentially expressed genes (DEGs) based on the authors’ annotations. We selected the cell context for each DEG extraction carefully to ensure that the resulting gene list accurately reflected true cell type or subtype markers. For instance, for cell subtype markers, we isolated cells within the context of their sibling cell types to pinpoint markers capable of distinguishing between subtypes.

We proceeded to verify and filter the curated marker gene sets by identifying markers that were consistently expressed across multiple datasets with the same author annotations. This step was crucial to exclude genes whose differential expression stemmed from factors other than cell type, such as age, gender, condition, or technology. Furthermore, we confirmed that these marker gene sets could identify distinct clusters in other datasets, regardless of differing nomenclature or broad-level annotations. This process facilitated the merging of different naming conventions under a unified cell population, subsequently labeled using the most appropriate terms from cell ontology from the European Bioinformatics Institute (EBI)^1^. In instances where our initial marker gene sets were derived from an unsuitable cell context - resulting in poor cross-study agreement - we revisited and refined the cell context selection for DEG extraction, repeating the process as necessary.

Upon refining markers for each cell type and subtype, we assessed their uniqueness across different tissues. When marker gene sets were unique to a cell type in certain datasets but ambiguous in others, we manually merged data from various tissues to establish a new context for identifying unique markers. This extensive curation effort led to the development of a comprehensive map comprising 166 cell type and subtype-specific marker gene sets, detailed in Supplementary Data 2. This map not only enhances the accuracy of cell type identification in scRNA-seq studies but also improves the reproducibility of research findings across the scientific community.

### Prediction algorithm

To automatically assign cell labels based on the curated marker gene sets, we developed a customized algorithm that builds upon the original AUCell algorithm^2^. In summary, there are three main steps: (1) Cut off the background noise: In scRNA-seq experiments, contamination and alignment errors can introduce false low-level gene expression. To address this, we first identify the expression threshold for each gene in the study and remove the lowly and randomly expressed genes. (2) Perform AUCell enrichment and assign cell labels: We apply the AUCell algorithm to calculate the enrichment score of each marker gene set. AUCell works by assessing the enrichment of predefined marker gene sets in the transcriptomic profile of each cell. We assign each cell a label corresponding to the marker gene set that shows the highest enrichment, thus effectively categorizing cells based on their gene expression profiles. (3) Smooth unassigned cells: Drop-out events - where a gene’s expression is not detected due to technical limitations rather than biological absence - are a common challenge in single-cell datasets. These events can lead to false negatives, where essential markers are not observed. To address this, our algorithm includes a smoothing step where unassigned cells are labeled based on the consensus of their neighboring cells. Specifically, we assign a label to an unassigned cell if its 15-nearest neighbors (in terms of gene expression profile similarity) have a consistent label. The detailed algorithm is described in the Method section.

We also implemented our prediction algorithm in a user-friendly application, which enables researchers to quickly and easily annotate their scRNA-seq data using our comprehensive marker gene sets. The application includes interactive visualizations and quality control metrics, as well as the ability to perform other downstream analyses. Overall, our prediction algorithm and application provide a powerful and accessible solution for accurate and efficient cell type annotation in scRNA-seq data.

**Figure 2.**
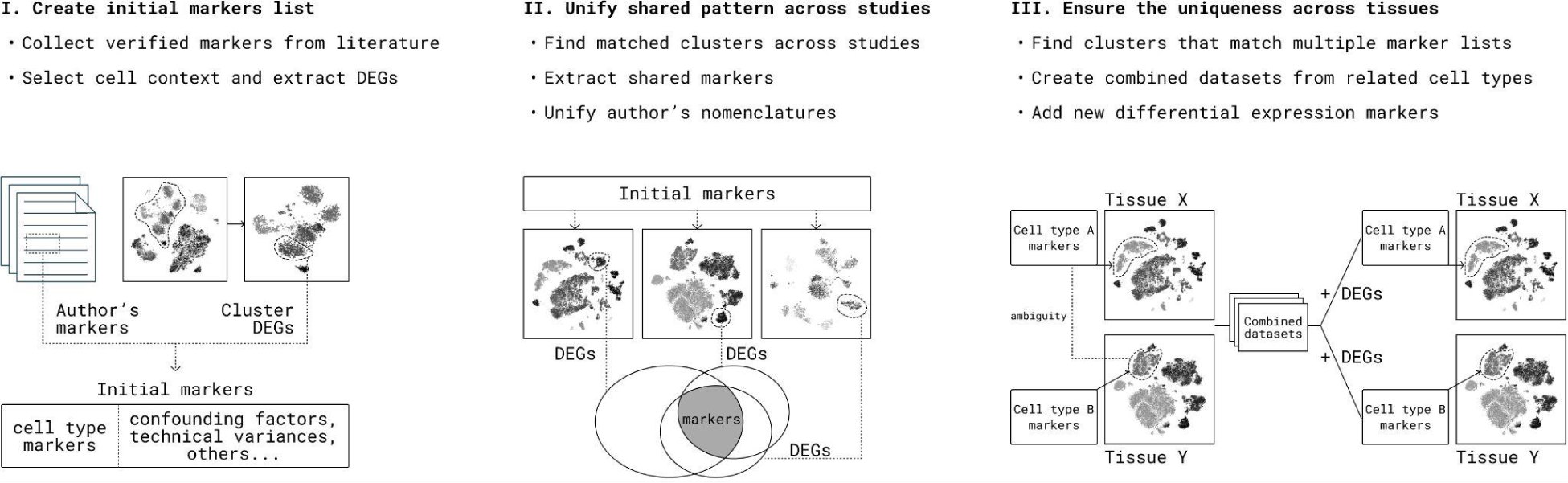
The process of curating marker gene sets for cell type annotation. The figure illustrates the three main steps involved in curating marker gene sets for accurate cell type annotation. (I) The initial step involves collecting verified markers from the literature and extracting differentially expressed genes using specific cell contexts. (II) The second step involves unifying shared patterns across different studies by finding matched clusters, extracting shared markers, and unifying nomenclature. (III) The final step involves verifying the marker’s uniqueness across different tissues. This step involves finding clusters that match multiple marker lists, combining datasets from related cell types, and adding new differential expression markers to refine and confirm the marker specificity.

### Benchmark results against reference-based mapping method

In evaluating the efficacy of our novel annotation tool, OmnibusX, we conducted comprehensive benchmarking against Azimuth, a widely used reference-based mapping method. Our benchmarking strategy involved selecting 22 datasets spanning 11 different tissues, including kidney, liver, lung, bone marrow, brain, heart, pancreas, peripheral blood, breast, colon, and eye (detailed in Supplementary Data 3). For each tissue, we chose two datasets: one from Azimuth’s reference collection and another external dataset not included in its reference. This approach allowed us to assess the performance of Azimuth in familiar and novel contexts, thereby testing its adaptability and accuracy in broader applications. For tissues such as peripheral blood where our initial annotation attempts using Azimuth’s reference did not succeed), breast, colon, and eye (where Azimuth’s references are not available), we utilized two datasets not included in Azimuth’s reference.

For each dataset, we performed cell type annotation using both our method and Azimuth, and then compared the results to the original author-provided cell type labels. Due to the different used terminologies by authors, Azimuth and OmnibusX, we then standardized the terminology across datasets using a controlled vocabulary, facilitated by cell ontology from the EBI. We used Cohen’s kappa coefficient^4^ to assess the general agreement between the predicted and true cell type labels, as it accounts for the possibility of agreement occurring by chance. Our results indicated that OmnibusX consistently outperformed Azimuth across most tissues (Figure 3A and Figure 3B), with higher Cohen’s Kappa scores in both reference and external datasets, achieving an average Cohen’s kappa coefficient of 0.88 across all datasets, while Azimuth achieved an average score of 0.75.

**Figure 3:**
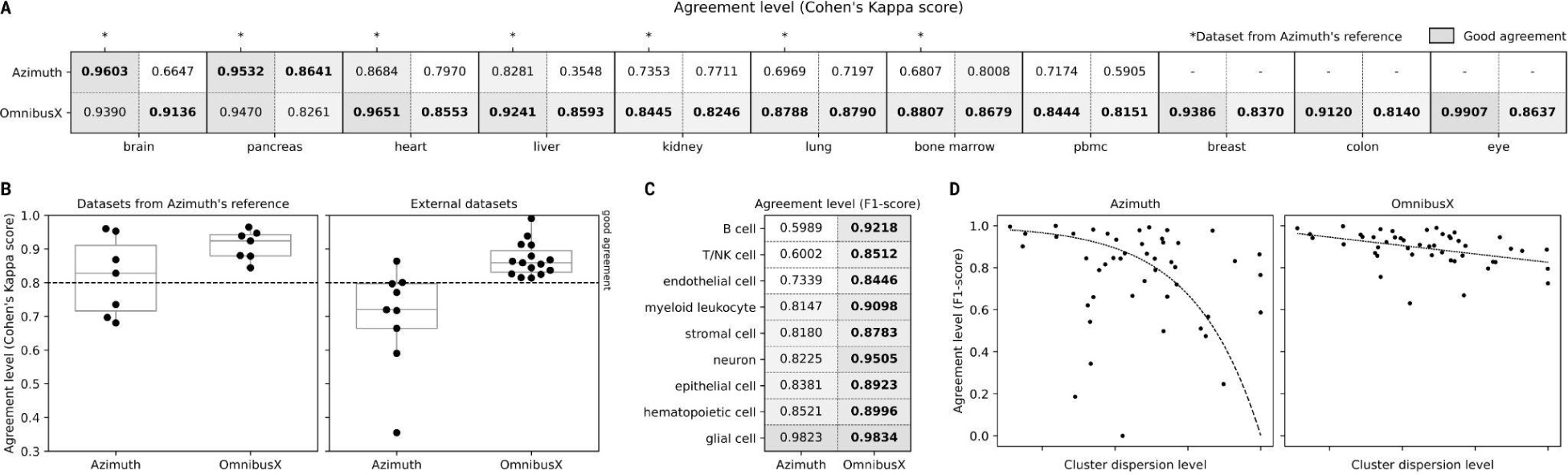
Comprehensive Analysis of Annotation Performance Between OmnibusX and Azimuth. **A.** Agreement levels (Cohen’s Kappa scores) by datasets between annotations made by OmnibusX and Azimuth compared to author annotations across various tissues. Scores marked with an asterisk indicate datasets from Azimuth’s reference. **B.** Box plots represent the Cohen’s Kappa scores for both tools between datasets from Azimuth’s references and external datasets, showing higher consistency for OmnibusX, especially on external datasets. **C.** Agreement level (F1-score) by cell types across 22 datasets, with OmnibusX generally outperforming Azimuth, particularly in subtypes with high cluster dispersion (B cell, T/NK cell). **D.** Correlation between the agreement level (F1-score) against the cluster dispersion level for both tools, highlighting Azimuth’s performance decline in highly dispersed clusters, whereas OmnibusX maintains a more stable performance across varying degrees of cluster dispersion.

We then combine the annotations from 22 studies and evaluate the performance of both tools on classifying subtypes of different cell types. Notably, OmnibusX maintained steady performance across different cell types, whereas Azimuth showed significant performance drops in classifying subtypes within B cell and T/NK cell categories (Figure 3C), which are often characterized by overlapping populations and dispersed clustering patterns. We then investigated the relationship between the performance of both tools and the degree of cluster dispersion, measured by the Silhouette Coefficient^5^. OmnibusX showed only a slight performance decline in more dispersed cell type clusters, while Azimuth’s performance decreased dramatically under similar conditions (Figure 3D). These findings are visually summarized in Figure 3, which illustrates the comparative analysis of agreement levels and performance metrics between OmnibusX and Azimuth across different datasets and cell types.

In summary, our benchmarking results affirm the robustness of the OmnibusX tool in providing reliable, consistent cell type annotations across a wide range of tissues, outperforming traditional reference-based methods, particularly in challenging scenarios involving dispersed clusters and closely related subtypes. The flexibility of our approach also allows for the inclusion of new cell subtypes as they are identified, enhancing the adaptability of our tool to new data. Our user-friendly application further facilitates the annotation process, significantly reducing the workload for researchers. Importantly, all benchmark predictions with OmnibusX were conducted using the OmnibusX desktop application on a standard desktop computer, demonstrating its scalability and ease of use for a wide range of scientists.

## Discussion

In this study, we present a novel method for cell type annotation in single-cell RNA sequencing data using a curated set of marker genes. Our approach addresses the challenges of inconsistent cell type nomenclature and the presence of closely related cell subtypes that are difficult to distinguish using clustering methods. By carefully selecting marker genes and verifying their specificity across multiple datasets and tissues, we have developed a comprehensive map of cell type and subtype-specific markers that can be used for accurate cell type annotation.

Our customized algorithm, which is based on the AUCell method, enables the automatic assignment of cell labels using our curated marker gene sets. We have benchmarked our method against a reference-based mapping tool called Azimuth and demonstrated that our approach consistently outperforms Azimuth in terms of agreement with author-provided cell type annotations. Importantly, our method is not limited to datasets included in the reference and can accurately annotate cells in external datasets with varied cell type composition. Furthermore, our method allows for the detection of closely related cell subtypes that may not be distinguishable using clustering methods or manual annotation. Another advantage of our approach is the use of a consistent marker gene set across all datasets, simplifying the process of cell type annotation and reducing the need for reference selection. Furthermore, our user-friendly application saves scientists time from coding and allows for easy verification of annotation results.

One potential limitation of our approach is the reliance on marker genes for cell type annotation. While we have carefully curated and verified our marker gene sets, it is possible that new markers may be discovered or that some markers may not be specific to certain cell types in all contexts. Therefore, it is important to continue updating and refining the marker gene sets as new data becomes available.

In conclusion, our method provides a powerful tool for accurate and consistent cell type annotation in single-cell RNA sequencing data. By addressing the challenges of inconsistent nomenclature and closely related cell subtypes, our approach enables more precise analysis of complex tissues and facilitates the discovery of new cell types and functions. The user-friendly application and comprehensive map of cell type and subtype-specific markers make our method accessible to a wide range of researchers in the field.

## Methods

### scRNA-seq data acquisition and processing

We obtained all scRNA-seq data used for verifying cell type and cell subtype markers from the Data Availability section of each publication. A complete list of these publications is provided in Supplementary Data 1. Raw counts were prioritized for download; however, if only normalized data were available, we utilized those and omitted the normalization step in our subsequent analyses.

For the benchmarking analysis, we downloaded data from multiple platforms: Scanpy objects were obtained from CellxGene^10^ (https://cellxgene.cziscience.com/) and the Human Cell Atlas (https://www.heartcellatlas.org/), and raw count matrices (.csv files) were downloaded from the Gene Expression Omnibus (GEO) and the Allen Brain Map (https://portal.brain-map.org/). We selected datasets from 11 different tissues, choosing two per tissue. For tissues with an existing Azimuth reference, one dataset was selected from the reference source to evaluate the performance of the reference-based tool across different data sources. The links to download each dataset can be found in Supplementary Data 3. For datasets received as Scanpy objects, we extracted the raw count matrices to facilitate subsequent analysis.

Data processing was performed using a standardized pipeline from Scanpy^7^. For the marker curation process, we adjusted the cell filtering threshold according to the authors’ descriptions and annotations, and cells without labels or with uncertain labels were removed. The raw count matrix was then normalized to a total count of 10,000 and log-transformed using the log1p function. We detected the top highly variable genes using default parameters (min_mean=0.0125, max_mean=3, min_disp=0.5) and applied Principal Component Analysis (PCA) and t-SNE using default parameters. No batch correction was implemented.

For the benchmarking analysis, we subset the original datasets to fewer than 50,000 cells per dataset to minimize the bias associated with large datasets when evaluating the performance of both tools across studies. Raw data were used directly for predictions using Azimuth, while the standard processing pipeline described above was applied for OmnibusX predictions. The results from t-SNE were utilized to assess the degree of cluster dispersion in subsequent sections. To establish a reference for later evaluation, we mapped the authors’ cell annotations to the corresponding standard terms from the Cell Ontology, which were then considered as the ground truth.

### The curation process for unique marker sets

1. Initial labeling and marker Identification: Cell type labels were primarily derived from the authors’ annotations in each study. In cases where direct annotations were unavailable, cell clusters were annotated based on markers mentioned in the study text. The initial list of marker genes was established using verified markers provided by the authors. Subsequently, for each cell type, we performed multiple rounds of t-tests (implemented by Scipy^8^) to extract additional differentially expressed genes (DEGs). These DEGs were selected based on strict criteria to minimize the impact of technical variance across datasets: only DEGs clearly distinguishing two groups with a p-value < 0.05, expressed in more than 15% of one group and less than 5% in the other, were retained.
2. Global and contextual marker refinement: Initially, each selected cell type was compared against all others within the same study to identify globally unique markers. This ‘global’ scope, however, is confined to the context of one study, significantly influenced by the cell type composition. For instance, if all immune cells are sorted from CD3+ cells, common immune markers like PTPRC could appear uniquely associated with T cells. Given these constraints, subsequent verification across multiple studies became essential. Generally, truly unique markers are rare, so we employed a multi-level contextual selection process to extract sets of genes that collectively distinguish the selected population. We categorized annotated cell types into broad compartments: epithelial, stromal, immune, neural, and stem/progenitor cells. DEG analysis was initially conducted between the compartment of the selected cell type and other compartments to derive a broad marker list. Further refinement was achieved by performing DEG analysis within the selected compartment, comparing the specific cell type against others to extract cell subtype markers. These two marker lists were then combined. Subsequent subsetting and DEG analysis are necessary for closely similar cell subtypes, which are evaluated and adjusted on a case-by-case basis.
3. Iterative verification and context adjustment: After establishing a preliminary marker list for a cell type from one study, we continued to verify and refine this list in subsequent studies featuring the same cell type. This process involved extracting markers for each study and intersecting them with existing lists to retain commonly shared markers. If the intersection significantly reduced the marker list, manual adjustments were made to address potential biases or mislabeling by authors. A marker list was deemed robust if it maintained consistency across at least ten different studies.
4. Definition of new cell types and subtypes: New cell types and subtypes, often identified in only a handful of studies, require meticulous evaluation to determine their inclusion or exclusion from the prediction list. Starting from the extract marker list from studies that define them, we programmatically scan through our available datasets for detecting cell populations enriched in the markers list (AUCell enrichment score > 0.3, irrespective of the author’s annotations). Studies were ranked by the number of matching cells, and each potential match was manually reviewed to confirm the presence of a consistent cell cluster. If confirmed in at least ten studies, including initial ones identifying the new type, the cell type, and its markers were added to our curation map.
5. Final marker list verification: Upon establishing the preliminary markers list for all detectable cell types, we methodically processed each study, applying AUCell enrichment analysis to all marker lists against each author’s cell type annotation. The enrichment score matching the author’s annotation served as the baseline threshold. Populations were identified as enriched in multiple marker lists if more than 30% of the cell population had enrichment scores for other markers exceeding this base threshold. To mitigate issues of overlapping due to background noise or doublets, we established another criterion: overlaps between two sets had to occur in more than 300 cells across at least two studies. Ambiguous marker lists underwent further refinement through manual consolidation of related studies, repetition of the DEG extraction process, and updates to the marker lists, ensuring the highest level of accuracy and reliability in our final cell type classifications. Each marker list was labeled using terms from the Cell Ontology.

All manual curation steps were conducted using the OmnibusX single-cell application. The programmatic processes were carried out on a Linux server utilizing Python scripts.

### Azimuth prediction

For the benchmark analysis, raw data from 16 selected datasets were uploaded to the Azimuth server (https://azimuth.hubmapconsortium.org/references/). Each dataset was aligned with the appropriate tissue-specific reference from Azimuth: brain (Human - Motor Cortex), pancreas (Human - Pancreas), heart (Human - Heart), liver (Human - Liver), kidney (Human - Kidney), lung (Human - Lung v2 (HLCA)), bone marrow (Human - Bone Marrow), blood (Human - PBMC). The result annotations were subsequently downloaded as a .tsv metadata file. In the case of the PBMC reference, we were unable to perform mapping annotations using Azimuth’s reference subset; thus, for this tissue, both datasets were external data.

Each reference provides different levels for mapping cell type annotations. We initially selected the level with the highest agreement with the author’s annotation. Cell subtypes that matched with the author’s annotation from a more granular level were manually merged to the selected level. Cells belonging to the same broader cell types were also manually merged to the broader name, in accordance with the author’s annotation level. Lastly, cell annotations were mapped to the standardized terms from the Cell Ontology.

### OmnibusX prediction methodology

1. Cut off the background noise: To minimize low-level signal interference, we set a cutoff threshold for each gene based on its expression values. Using raw counts as input, the expression threshold for each gene is determined by the 90th quantile of expression values among cells that express that gene. If a gene’s expression falls below this threshold divided by 10, it is considered noise and is removed from the analysis.
2. Perform AUCell enrichment and assign cell labels: AUCell enrichment analysis is conducted for every marker set across all cells. The enrichment threshold for each cell is defined as the greater of 0.1 or (max_score - 0.1), where max_score represents the highest AUCell enrichment score achieved by any marker set within that particular cell. Cells enriched in multiple marker sets are assigned to the set with the greater number of expressed genes (expression count > 0). Cells that do not match any marker set or express fewer than half of the genes in any given marker set are labeled as ‘Unassigned’.
3. Smooth unassigned cells: Using PCA results from the prior processing step, we calculate the 15-nearest neighbors for each cell using PyNNDescent^11^. For cells labeled as ‘Unassigned’, we examine their neighbors. If no label occurs at least twice among the neighbors, the cell remains ‘Unassigned’. For each label that appears in a minimum of two neighbors, we calculate the average distance of the cells in each label to the unassigned cell and assign it to the label of the closest group.

Steps 2 and 3 were repeated twice. The first iteration was performed using broad cell type marker lists, followed by a second iteration for each broad cell type using their respective subtype marker lists. This process resulted in two levels of cell type annotation: broad cell types and subtypes. By performing the process consecutively, OmnibusX ensures consistency between the two levels of annotation, providing an advantage over the current Azimuth strategy. Azimuth maps each cell type level independently, which can lead to discrepancies between level annotations, such as a T cell subtype being identified as a naive B cell. This discrepancy also requires significant time to resolve inconsistencies.

For the purpose of benchmark evaluation, we also merge the OmnibusX prediction subtypes to broader cell type labels, in accordance with the author’s annotation level.

### Benchmark metrics

To assess the performance of each tool across different tissues, we utilized Cohen’s Kappa scores, calculated using the Scikit-learn package, with the authors’ annotations serving as the ground truth. The Kappa score quantifies the agreement between two annotations; it ranges from −1 (no agreement) to 1 (perfect agreement). Scores above 0.8 typically indicate good agreement, while scores at or below zero suggest no agreement, akin to random labeling. We also presented the scores as box plots, illustrating the score distribution across tissues from both Azimuth’s reference datasets and external datasets. This comparison highlighted the consistent performance of OmnibusX across datasets, while Azimuth’s performance was notably lower on external datasets, as anticipated.

Upon examining the prediction results, we noticed a performance bias associated with the dispersion of cell types. Consequently, we further evaluated the performance of both tools against different cell types, characterized by varying degrees of dispersion across tissues. We combined the prediction results from all 16 datasets that Azimuth could predict and categorized the cell type labels into different groups of broad cell types. These groups included: B cell, T/NK cell, epithelial cell, glial cell, endothelial cell, myeloid leukocyte, hematopoietic cell, stromal cell, and neuron. We then calculated the subtype label call accuracy of each tool for each broad group using the F1-score (Scikit-learn package). Due to the loss of context from each study assignment when extracting and combining subsets of cell types, Cohen’s Kappa score was not suitable for this evaluation. As expected, Azimuth’s performance on closely related subtypes, such as T/NK cells and B cells, dropped significantly.

We further investigated the correlation between cell type dispersion and the performance of each tool using t-SNE coordinates from the processed data. For this analysis, each study’s cell type annotations were divided into the aforementioned broad groups. We then calculated the Silhouette Coefficient (Scikit-learn package) for the subtype labels within each group using t-SNE coordinates. The Silhouette Coefficient, which ranges from −1 (incorrect clustering) to 1 (perfect clustering), assesses the clustering quality based on the mean intra-cluster distance (a) and the mean nearest-cluster distance (b). Values near 0 indicate overlapping clusters, suggesting ambiguity in cluster assignments. Subsequently, we computed the F1-scores for subtype labeling for both tools. Data points representing the pair of Silhouette Coefficient scores and F1-scores were plotted for each broad cell type group combined across 16 studies. Broad cell types with no subtype annotation from the authors or containing less than 1000 cells were skipped to reduce random scores due to small sample size. To provide a clearer visualization of trends, we reversed the Silhouette Coefficient scale, illustrating performance decay from left to right. This approach revealed an exponential decline in Azimuth’s performance with increasing dispersion, whereas OmnibusX showed only a slight decline. All plots were made using Matplotlib^9^ and Seaborn^12^.

## Supporting information

Supplementary Data 1: List of publications referenced for the curation process

Supplementary Data 2: List of predictable cell types and subtypes

Supplementary Data 3: Benchmark datasets

## Data availability

All relevant data supporting the key findings of this study are available within the article and its Supplementary Information files. All data used as part of this work is publicly available from the cited studies. Data subsets used for benchmarking, along with source data to recreate figures, have been deposited on Zenodo: https://doi.org/10.5281/zenodo.11183242. Additionally, a selection of curated marker genes for cell types and subtypes is available from the corresponding author upon reasonable request.

## Code availability

OmnibusX is accessible as a closed-source application at https://omnibusx.com. All code necessary to reproduce the benchmark results is available at https://github.com/OmnibusXLab/celltype-prediction-benchmark.

## Acknowledgments

We thank OmnibusX Company Limited for supporting this research and providing access to the resources necessary to complete the project. Although the code used in this study is not publicly available due to proprietary reasons, we are willing to discuss the methods and results with interested researchers to the extent allowed by our confidentiality agreements.

## Author contributions

Conceptualization, methodology, data curation and analysis, manuscript writing and coding: Linh Truong, Thao Truong, and Huy Nguyen.

## Additional information

**Supplementary information** Supplementary Information is available for this paper.

Correspondence and requests for materials should be addressed to Huy Nguyen at.

